# Examining Intercage Transmission of *Chlamydia muridarum*: Impact of Barrier Husbandry and Cage Sanitization

**DOI:** 10.1101/2024.04.19.590256

**Authors:** Michael B Palillo, Noah Mishkin, Mert Aydin, Anthony Mourino, Rodolfo J Ricart Arbona, Neil S Lipman

## Abstract

*Chlamydia muridarum* (Cm) has reemerged as a prevalent bacterial contaminant of academic research mouse colonies. A study was conducted to assess the effectiveness of husbandry and cage sanitization methods in preventing intercage transmission of Cm. To assess intercage transmission during cage change, a cage housing 2 Cm-free Swiss Webster (Tac:SW; SW) sentinel mice was placed randomly on each of 12 individually ventilated cage racks, housing cages with Cm-shedding mice, located in 1 of 2 animal holding rooms. Husbandry staff blinded to the study cages, changed all cages in the animal holding rooms weekly using microisolator cage technique. PCR testing performed 180 days post-placement confirmed all mice remained negative for Cm. To assess the effectiveness of cage sanitization to eliminate Cm, we investigated transmission of Cm to a naïve Cm-free SW and NOD.Cg-*Prkdc*^scid^ *Il2rg*^*tm1Wjl*^/SzJ (NSG) mouse co-housed for 7 days (repeated weekly for 4 weeks) in cages assigned to 1 of 3 groups (n=10 pairs of mice/group). Cages that previously housed 2 Cm-shedding BALB/c mice were either washed in a tunnel washer (82.2°C [180°F] final rinse for an average of 16 seconds per run; n=10) with and without post-washing autoclaving (121°C for 20 minutes; n=10), or were untreated (bedding change only; n=10). Pre- and post-sanitization swabs of each cage were assayed for Cm by PCR. All pre-treatment swabs tested positive, while post-treatment swabs from all cages (excluding bedding change) tested negative. All SW and NSG mice, irrespective of group, remained negative for Cm as determined by PCR. These findings suggest that infectious Cm does not persist in untreated cages nor after mechanical washing with and without autoclaving. Collectively, these findings suggest that neither our husbandry protocols nor inadequate cage sanitization methods likely contributed to the observed prevalence of Cm in contemporary research mouse colonies.

## Introduction

*Chlamydia muridarum* (Cm) is a gram-negative, obligate, intracellular bacterial pathogen in the Chlamydiaceae family and the sole known natural chlamydial pathogen of mice.^5,26^ Discovered in the 1930s in association with respiratory disease and lung pathology in research mice, Cm has subsequently been widely utilized as a model for investigating human *Chlamydia trachomatis* infection.^13,18,31^ Despite its widespread experimental use, the biology of natural Cm infection remains relatively uncharacterized, with limited information available on its infectivity and environmental stability.^15,31^

Cm was recently isolated from and shown to be prevalent and globally distributed among research mouse colonies.^26^ Of 58 groups of mice imported from 39 academic institutions, 33% tested positive for Cm upon arrival, and 16% of a 900-diagnostic sample cohort and 14% of 120 institutions submitting approximately 11,000 murine microbiome samples had evidence of Cm.^26^ Cm infection, through its presumptive fecal-oral route of transmission of its metabolically inactive extracellular infectious elementary bodies (EBs), has been shown to cause significant disease, including severe pneumonia, dyspnea, and mortality, in highly immunodeficient and less severe pulmonary and potentially renal lesions in genetically engineered mouse (GEM) strains with select immune system alterations.^24,25,26,35^ Given Cm’s ability to induce a robust immune response, cause pathology in immunocompromised mice and persistent infection in both immunocompromised and immunocompetent mice, Cm should be excluded from research mouse colonies.^24,25,26,35^

Cm is also highly prevalent within our program where 62.9% of the mouse holding rooms house Cm infected mice.^26^ Its notable inter and intrainstitutional prevalence begs the question as to how the bacterium, which had not been detected in laboratory mice for almost a century, went undetected and became so widespread.^26^ One potential explanation for its distribution may be attributed to the massive global sharing of unique GEM strains among numerous research mouse colonies during the last 3 decades, as the method of introducing most of these mice into established colonies consisted of quarantine and testing for excluded agents. As Cm’s presence was unrecognized, and as a result no available commercial diagnostic testing was available until recently, the bacterium was presumably introduced inadvertently.

Even if the aforementioned method led to its’ interinstitutional spread, the high intrainstitutional prevalence leads to speculation that Cm may easily spread horizontally via aerosols and/or fomites. At most research institutions, GEM strains are housed in individually ventilated microisolator cages (IVC) which are only manipulated within specially designed ventilated animal changing/transfer stations or biological safety cabinets using handling techniques that are collectively designed to prevent spread of infective agents between cages. Could Cm’s prevalence be attributable to the failure of these husbandry methods as was demonstrated with the murine bacterial pathogen, *Corynebacterium bovis*?^6^ While many institutions autoclave IVC components between use, others depend on the effectiveness of heat and/or chemical sanitization provided by mechanical cage washing equipment.^7,32^ Therefore, alternatively or in addition to the ease of horizontal transmission, could inadequate cage sanitization lead to the persistence of Cm’s infective EBs on cage surfaces facilitating intercage transmission despite the use of the state-of-the-art caging and husbandry techniques? The studies described herein were devised and conducted to better understand whether either or both these mechanisms could be a factor in the spread of Cm within mouse colonies.

## Materials and Methods

### Experimental design

#### Intercage transmission study

A longitudinal study utilizing naïve, Cm-negative (confirmed via PCR) Swiss Webster (SW) mice was conducted with the primary objective of assessing the potential for intercage transmission of Cm resulting from routine barrier husbandry practices. Over 180-days, 12 cages each containing two naïve Cm-free female SW mice were placed on 12 Cm positive racks (1 cage/rack) in two mouse housing rooms located in separate barrier facilities. These 12 racks housed approximately 700 cages in total. The animal housing rooms and racks were selected based on their recently documented high prevalence of Cm-positive mice as detected using soiled bedding sentinel mice. Each cage of SW mice was randomly placed into an available spot on its designated rack. Over the 180-day period, cages were changed weekly by animal care technicians blinded to the study. All technicians adhered to the institution’s standard operating procedures for barrier cage change. To enhance the likelihood of changing a Cm-positive mouse cage before those added for the study, the study cages were randomly relocated to other available spots on the rack every 2 to 4 weeks throughout the study’s duration.

Feces was collected from each pair of SW mice (direct collection, pooled per cage) on the day of initial cage placement (day 0) and then on day 14, 30, 90, and 180. These samples were stored at −80°C (−112°F) until tested for Cm by PCR. After the final fecal collection, all study racks were re-tested for their Cm status. On day 180 after cage placement, mice were euthanized by carbon dioxide asphyxiation in accordance with recommendations in the AVMA Guidelines on Euthanasia of Animals, 2020 edition.^2^

#### Cage wash study

A randomized controlled trial employing mice shedding Cm was undertaken to investigate the efficacy of various cage sanitization methods in mitigating the spread of Cm within laboratory settings. Cages (n=30) each housing 2 female BALB/c mice shedding Cm (confirmed by PCR), changed weekly, were used to generate contaminated cages (n = 120). After soiled bedding was removed from the contaminated cage, a cotton applicator (Daiso, Hiroshima, Japan) was used to swab the interior perimeter of the cage bottom and then from corner to corner in an “X” pattern. Swabs were stored at −80°C (−112°F) before conducting PCR on an extract from each swab. Each contaminated cage was then randomly assigned to 1 of 3 groups: 1) WA: sanitization in a tunnel washer followed by autoclaving (n = 40); 2) TW: sanitization in a tunnel washer only (n = 40); or, 3) BR: bedding removed with no further cage sanitization (n = 40). The study design is illustrated in Figure 1A.

**Figure 1.**
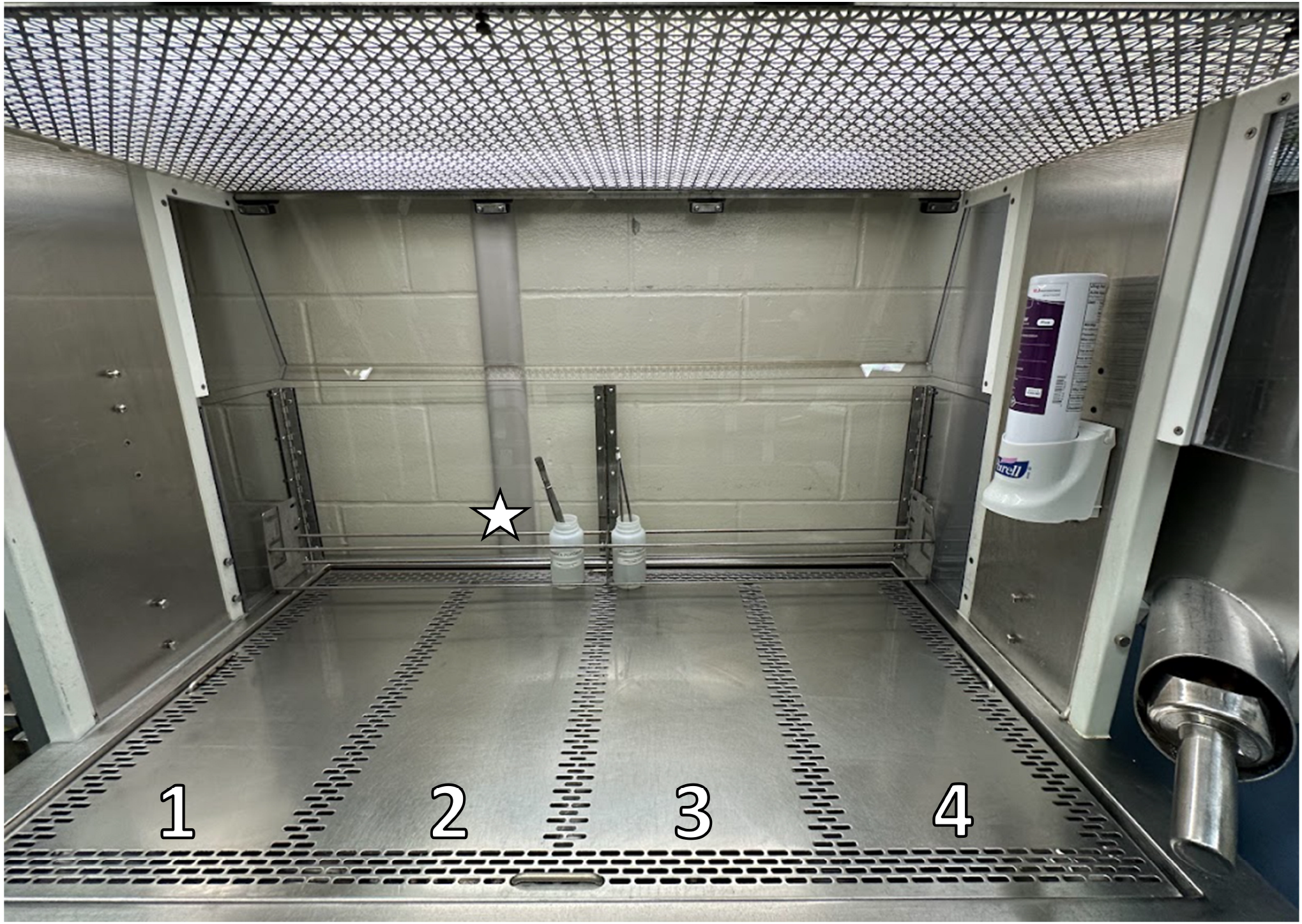
Schematic of the experimental design, including: (A) sampling and treatment of soiled caging and (B) timeline of exposure of SW/NSG pairs to treated test cages. The schematic represents housing of a single pair of mice from treatment groups for a total of four 1-wk periods.

Cages assigned to the BR group were not sanitized further. Autoclaved bedding, enrichment, food, water, wire bar lid (WBL), and filter top, as described below, were added to each BR cage along with 1 female SW mouse and 1 female NSG mouse. After soiled bedding removal, cages assigned to the WA and TW groups were sanitized without chemicals in a tunnel washer (Basil 6300, Steris, Mentor, OH) operating with a belt speed of 6 linear ft/min (1.8 m/min). A data logger (OM-CP-HITemp140, Omega Engineering, Bridgeport, NJ) was secured to the interior bottom of a nonexperimental cage and processed through the tunnel washer to confirm temperatures as the cage progressed through the washer immediately before the experimental cages were washed (Figure 2). The washer’s sump set points were as follows: 40-s cold water prewash, 40-s wash at 88°C (190°F), 30-s rinse at 88°C (190°F), 20-s final rinse at 88°C (190°F), and drying with an air knife system at 2,200 cubic feet per minute (128 m/s) air flow at 88°C (190°F) for 1 min. The maximum temperature to which the cages were subjected was confirmed using temperature-sensitive tape (Thermostrip, Cole-Parmer, Vernon Hills, IL) placed on the flat surface of a wire-bar lid immediately before the experimental cages were processed through the tunnel washer. Real time operational parameters displayed on the washer’s operator interface screen were also evaluated during cage processing.

**Figure 2.**
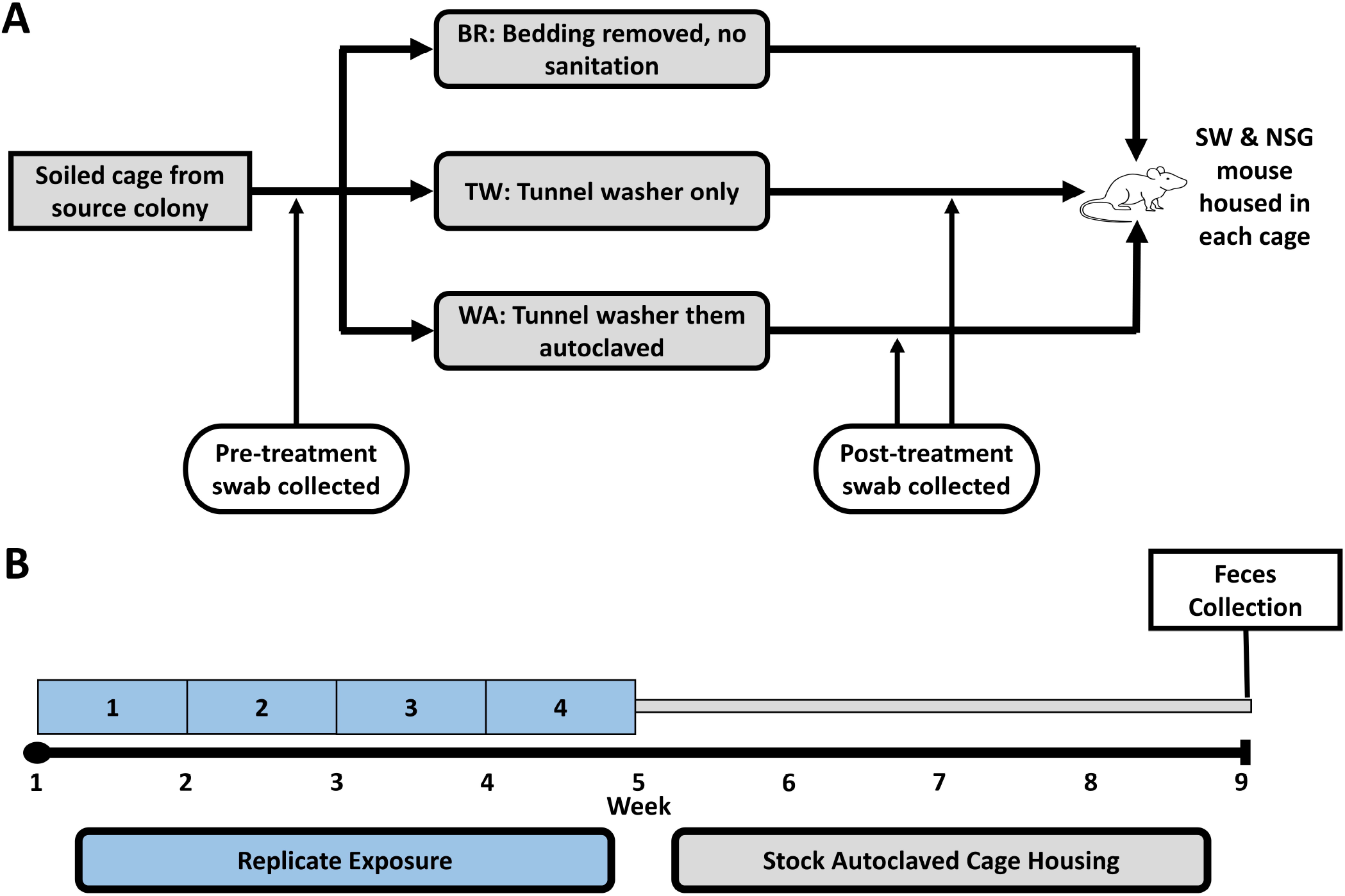
Temperature recorded by data logger run through the tunnel washer (N=4 runs). Data presented as mean ± SD.

After washing and retrieval from the tunnel washer, cages in the TW group were placed in a Class II, type A2 biologic safety cabinet (BSC; LabGard S602-500, Nuaire, Plymouth, MN) and swabbed for Cm as described above. WA group cages were fitted with an autoclaved filtertop after retrieval from the tunnel washer. The filtertop was secured in place with heat sensitive autoclave tape (Medline, Mundelein, IL), and the cage was autoclaved (Century SLH Scientific, Steris, Mentor, OH). After autoclaving, cages in the WA group were placed in the BSC and swabbed for Cm. After swabbing, autoclaved bedding, enrichment, wire bar lid, food, and water, processed as described below were placed in each TW and WA group cage, followed by the addition of a female SW and NSG mouse. For the entirety of the project, mice were housed in an isolation cubicle dedicated to this study. The cage wash and housing processes for each of the 3 groups were repeated so that each of the 10 pairs of mice was housed in a different cage from the same treatment group for four 1-wk exposure periods, resulting in testing a total of 120 contaminated cages (40 cages/treatment group). The time interval from removal of soiled bedding to placement of mice was 12-14 h for TW and WA cages. After the final 1-wk exposure period, mice were housed in autoclaved cages for the remainder of the study (Figure 1B). Four weeks following the last exposure period, fecal samples were collected directly from each mouse and pooled by cage. The fecal samples were stored at −80□°C (−112 °F) until tested for Cm by PCR. Mice were then euthanized by carbon dioxide asphyxiation.^2^

### Animals

Four- to 7-wk-old female SW mice (*n* = 24; Tac:SW, Taconic Biosciences, Germantown, NY) were used in the *intercage transmission study*. Three- to 6-wk-old female SW mice (*n* = 15) and 3-to 6-wk old female NOD.Cg-*Prkdc*^*scid*^*Il2rg*^*tm1Wjl*^/SzJ (n=15; NSG, Jackson Laboratories, Bar Harbor, ME) were used in the *cage wash study*. All mice were individually ear-tagged, assigned a distinct numerical identifier, and randomly assigned to appropriate groups (https://www.random.org/lists/) as described above. Cm-shedding female BALB/cJ (C) mice (*n* = 30; age 42 to 49-wk-old; The Jackson Laboratory, Bar Harbor, ME) were used to generate contaminated cages for the *cage wash study*. These mice were orally inoculated with a wild-type stock (Cm field strain isolated from clinically affected NSG mice at MSK) at a concentration of 2.72×10^4^ IFU in 100 uL of sucrose-phosphate-glutamic acid buffer (SPG, pH 7.2) using a sterile 18 g or 20 g orogastric gavage needle 7 months earlier. These mice were confirmed to be shedding Cm via fecal PCR on the first day of the study.

All mice obtained from vendors were free of mouse hepatitis virus (MHV), Sendai virus, mouse parvovirus (MPV), minute virus of mice (MVM), murine norovirus (MNV), murine astrovirus 2 (MuAstV2), pneumonia virus of mice (PVM), Theiler meningoencephalitis virus (TMEV), epizootic diarrhea of infant mice (mouse rotavirus, EDIM), ectromelia virus, reovirus type 3, lymphocytic choriomeningitis virus (LCMV), K virus, mouse adenovirus 1 and 2 (MAD 1/2), polyoma virus, murine cytomegalovirus (MCMV), mouse thymic virus (MTV), hantaan virus, murine chaphamaparvovirus-1(MuCPV), *Mycoplasma pulmonis*, CAR bacillus, *Chlamydia muridarum, Citrobacter rodentium, Rodentibacter pneumotropicus, Helicobacter spp*., *Salmonella spp*., *Streptococcus pneumoniae*, Beta-hemolytic *Streptococcus spp*., *Streptobacillus moniliformis, Filobacterium rodentium, Clostridium piliforme, Corynebacterium bovis, Corynebacterium kutscheri, Staphylococcus aureus, Klebsiella pneumoniae, Klebsiella oxytoca, Pseudomonas aeruginosa*, fur mites (*Myobia musculi, Myocoptes musculinis*, and *Radfordia affinis*), pinworms (*Syphacia spp*. and *Aspiculuris spp*.), *Demodex musculi, Pneumocystis spp, Giardia muris, Spironucleus muris, Entamoeba muris, Tritrichomonas muris*, and *Encephalitozoon cuniculi* when studies were initiated unless otherwise noted.

### Husbandry and housing

Mice were maintained in individually ventilated polysulfone cages with stainless-steel WBLs and filter tops (# 19, Thoren Caging Systems, Inc., Hazelton, PA) on aspen chip bedding (PWI Industries, Quebec, Canada) at a density of 2 mice per cage. Each cage was provided with a bag constructed of Glatfelter paper containing 6 g of crinkled paper strips (EnviroPak^®^, WF Fisher and Son, Branchburg, NJ) and a 2-inch square of pulped virgin cotton fiber (Nestlet^®^, Ancare, Bellmore, NY) for enrichment. Mice in the *intercage transmission study* were fed a closed source, natural ingredient, flash-autoclaved, λ-irradiated diet (experimental and colony; LabDiet 5053, PMI, St. Louis, MO), while mice in the *cage wash study* were fed a natural ingredient, closed source, gamma irradiated, autoclavable diet (5KA1, LabDiet, Richmond, VA). All mice were provided reverse osmosis acidified (pH 2.5 to 2.8 with hydrochloric acid) water in polyphenylsulfone bottles with stainless-steel caps and sipper tubes (Tecniplast, West Chester, PA) ad libitum. Cages in the *intercage transmission study* were changed every 7 d within an animal changing station (ACS, NU-S612-400, Nuaire, Plymouth MN), while cages in the *cage wash study* were changed every 7d within a class II, type A2 biological safety cabinet (LabGard S602-500, Nuaire, Plymouth, Mn). The rooms were maintained on a 12:12-h light:dark cycle, relative humidity of 30% to 70%, and room temperature of 72 ± 2°F (22 ± 1°C).

For the *cage wash study* only, select caging was autoclaved using a pulsed vacuum cycle of 4 pulses at a maximum pressure of 12.0 psig (6.9 kPa), with sterilization temperature of 121°C (250°F) for 20 min, and a 10.0 in Hg vacuum dry (3.4 kPa). Sterilization was confirmed by autoclave tape color change and post hoc verification of cycle chamber operating conditions. In addition, autoclave performance was verified weekly by using biologic indicators (Attest Biologic Indicators, 3M, Saint Paul, MN). Water bottles were autoclaved at a temperature of 121°C (250 °F) for 45 min with a purge time of 10 min.

The animal care and use program at Memorial Sloan Kettering Cancer Center (MSK) is accredited by AAALAC International, and all animals are maintained in accordance with the recommendations provided in the Guide.^22^ All animal use described in this investigation was approved by MSK’s IACUC in agreement with AALAS position statements on the Humane Care and Use of Laboratory Animals and Alleviating Pain and Distress in Laboratory Animals.

### Cage change

Animal care staff don dedicated vivarium scrubs and shoes and a disposable bouffant cap before entering the barrier. Barrier entry includes processing of shoes in a motor driven shoe cleaner (Shoe Brush 2001-TB, Liberty Industries, East Berlin, CT), stepping onto an adhesive mat (24”x36” White Tacky Mat, American Protective Products, North Branford, MA) and then proceeding through an air shower (AS2, Liberty Industries Inc, East Berlin, CT) into the barrier. Immediately following entry, disposable gloves are donned and treated with alcohol foam (Purell Advanced Instant Hand Sanitizer, 4121-06, GOJO Industries Inc. Akron, OH).

The cage change process is designed to prevent cross-contamination of infective agents between cages during a four-week cage change cycle. This is achieved by minimizing direct gloved hands contact with the interior of the cage, sanitizing gloved hands when transitioning between the cage’s inner and outer surfaces, and utilizing forceps soaked in a chlorine dioxide disinfectant (Clidox 1:18:1; Pharmacal Research Laboratories, Waterbury, CT) to handle animals and enrichment materials. Prior to initiating cage change, staff don a disposable gown and bonnet.

The first week of the cage change cycle involves a complete cage change, accompanied by the introduction of a Glatfelter paper bag containing 6 g of crinkled paper strips (EnviroPak^®^, WF Fisher and Son Inc) for enrichment. During the second week, the soiled cage bottom and bedding are replaced, the enrichment is transferred to the clean cage, and a 1” compressed cotton square (Nestlet^®^, Ancare Corporation) is added. This cage change includes replenishing feed levels and replacing water bottles, as needed, along with the transfer of the existing WBL and filter top. The next weekly change is a complete cage change with the transfer of existing enrichment items. During the fourth and final cage change of the cycle, the cage bottom and bedding are once again replaced with the transfer of the existing enrichment materials replenishing feed levels and water bottles, as necessary.

Cage change is conducted within an ACS (NU-S612-400, Nuaire) equipped with an external food hopper and scoop located on the ACS’s right side and its work surface is divided into four quadrants, numbered 1-4 from left to right (Figure 3). Before initiating cage change, the ACS’s interior surfaces are meticulously cleaned using a chlorine dioxide-saturated cloth (WypAll Economizer L30 ¼ Fold Wipers, Kimberly-Clark, Roswell, GA) and the ACS is turned on allowing air flow to stabilize. Following donning and sanitizing new gloves with an alcohol-based foam (Quik-Care Aerosol Foam Hand Sanitizer, ECOLAB, Saint Paul, MN), two forceps jars containing large blunt tipped forceps (VWR^®^ 10” Serrated Specimen Forceps, Avantor, Radnor Township, PA) filled with chlorine dioxide solution (Clidox 1:18:1; Pharmacal Research Laboratories) are placed in a holder at the rear of the ACS (Figure 3). An empty cage bottom, to be used for housing soiled bedding sentinel mice, containing a disposable plastic sample cup (1oz Plastic Medicine Cups, Owens & Minor Inc., Mechanicsville, VA) is placed in section 1. The ACS is organized, and the cage change process is performed such that the sanitized clean cage changing materials are placed on the ACS’s right side while the soiled materials on the left.

**Figure 3.**
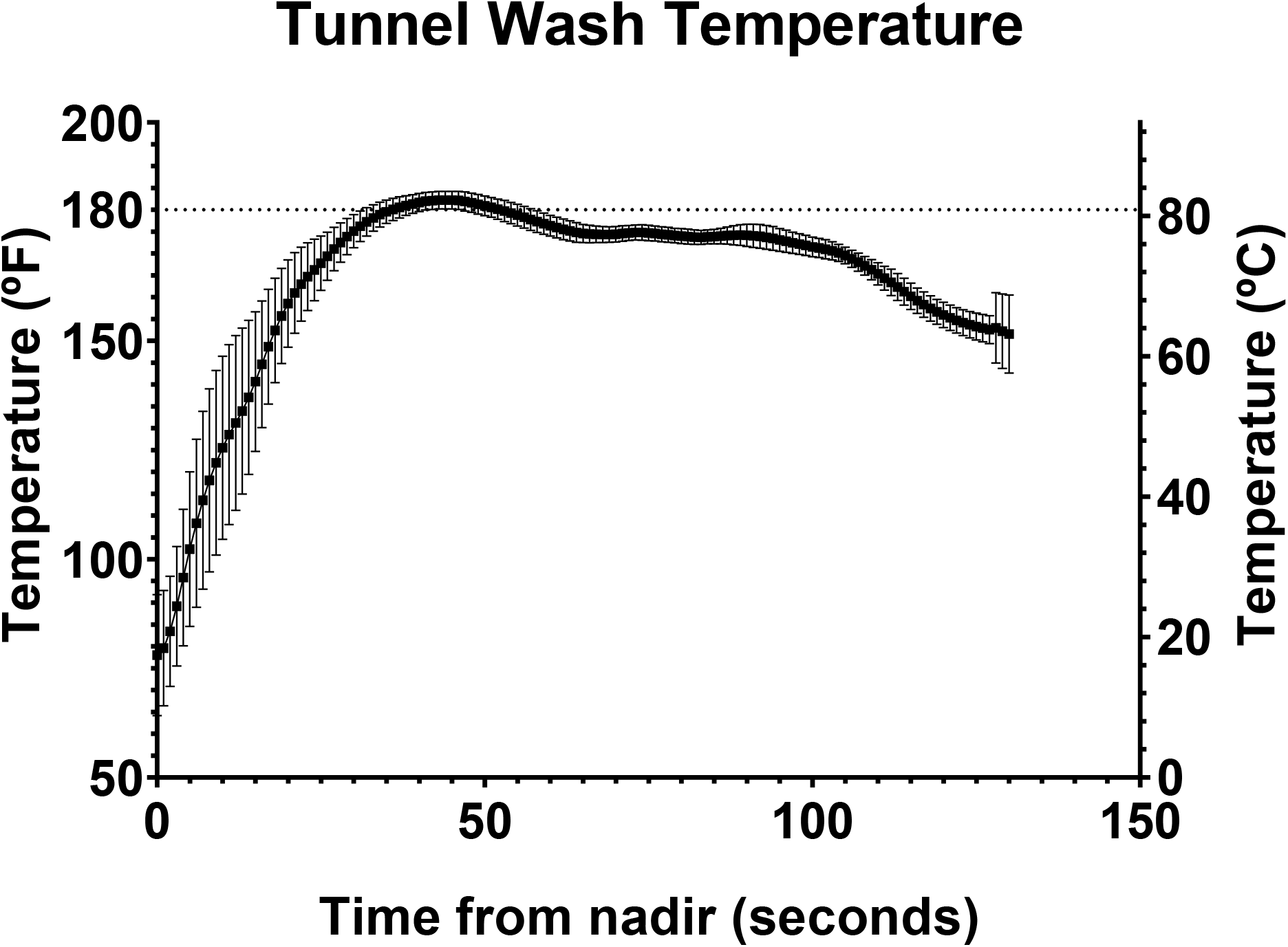
ACS with the four respective quadrants, numbered 1-4 from left to right. (1) Position for empty sentinel cage & disposable plastic cup; (2) Position for soiled cage; (3) Position for clean cage; (4) Position for face up filter top during complete cage change; and, ACS holder (used during bottom cage change) with forceps jars filled with chlorine dioxide solution containing large blunt tipped forceps (star).

Cage change begins with the removal of the upper right-hand cage of the rack to be changed proceeding horizontally from right to left following a reverse-S pattern. When conducting a complete cage change, the soiled cage is positioned in section 2 of the ACS. Subsequently, a clean cage setup is placed in section 3 of the ACS and its’ filter top is removed, with contact limited to the top’s outer edge, which is then positioned face up in section 4 of the ACS. The filter top is then removed from the soiled cage ensuring contact is limited its’ outer edge and it is leaned against the stainless crossbar in section 2 of the ACS. Gloved hands are then again moistened with foam and the clean WBL is removed and placed on the inverted filter top in section 4. The soiled water bottle is removed from the WBL and placed into a soiled water basket on a cart located adjacent to the ACS. The soiled WBL is lifted and is rested on the outer edge of the soiled cage using the non-animal handling hand. Forceps that have been immersed in the chlorine dioxide solution for the longest duration are used to transfer the existing enrichment materials and animals. The forceps are then replaced in the forceps jar to ensure adequate contact time between uses. The cage card holder attached to the front of soiled cage is then transferred to the clean cage. The clean WBL is replaced, and a clean water bottle is added. The food remaining from the soiled WBL is carefully poured into the clean WBL’s food container. Hands are again moistened with foam and feed is topped off using a food scoop, taking care to avoid touching the WBL or the existing food. The clean filter top is replaced, handling only its outer edge, followed by replacement of the soiled filter on the respective cage. The changed cage is returned to its original position and the process repeated for every cage on the rack side. Upon completion of cage change for all cages on a rack side, the chlorine dioxide solution in the forceps jars is replaced, the interior surfaces of the ACS are again cleaned using a chlorine dioxide-saturated cloth and new gloves are donned.

During the second and fourth week of the cage change cycle, a bottom only cage change is performed consisting of replacement of the soiled cage bottom and bedding in addition to feed replenishment and replacement of water bottles, as needed. The preparatory steps, organization of materials, and the cage change process are repeated as described above for the complete cage change with select differences. Following the placement of a clean bedded cage in section 2 and placement of the soiled cage to be changed in section 3, the filter top is carefully removed from the soiled cage, ensuring contact is limited to its’ outer edge, and it is leaned against the stainless crossbar in section 2 of the ACS (Figure 3). Gloved hands are then re-moistened with foam and the existing WBL on the cage to be changed is lifted and rested upon the outer rim of the soiled cage, using the non-animal handling hand. Forceps that have been immersed in the chlorine dioxide solution for the longest duration are used to transfer the existing enrichment materials and animals into the clean cage bottom. The existing WBL is placed onto the clean cage. Hands are moistened with foam and feed is carefully replenished as described above and the existing filter top is then replaced.

Weekly, one column of cages is designated for inclusion in the soiled bedding sentinel colony health monitoring program. When changing cages in the identified column, ∼15mL of bedding from the soiled cage is collected and placed into the empty cage in section 1 using the disposable plastic sample cup prior to replacing the filter top (Figure 3). The cup is left in the sentinel cage until all cages undergo the change. Lipman and Homberger have previously described the specifics of sentinel animal housing and the soiled bedding sentinel colony health monitoring program.^23^

### Cm positive holding room and rack selection

The 2 rooms used in the *intercage transmission study* were selected based on their documented high prevalence of Cm based on the soiled bedding sentinel colony health monitoring program’s findings. These rooms collectively contained 11 single-sided and 7 double-sided rack units (Thoren Caging Systems, Inc.). A “rack” is defined as a single-sided rack unit or one side of a double-sided rack unit, each rack holding up to 70 cages. To identify specific racks per room (6 per room, 12 total in study) to be used in the study, approximately 15 mL of soiled bedding was collected from each cage present on each of the 25 racks and placed into an autoclaved cage. Two sterile cotton-tipped applicators (Daiso, Hiroshima, Japan) were inserted into each of the 25 cages generated and an autoclaved filter top was affixed to the cage. The cage was then shaken 20 times horizontally (left to right to left) and 20 times vertically (up to down to up) to ensure full exposure of the swabs to the soiled bedding. This process was replicated for each of the 25 racks, and the swabs were stored at −80°C (−112°F) until tested for Cm via PCR. Twenty-one of the 25 racks were determined to house Cm-shedding mice and 12 racks were randomly selected for study inclusion (https://www.random.org/lists/). This process was repeated at the end of the study to confirm the continued presence of Cm-shedding mice on each of 12 racks used in the study.

### Fecal collection

Fecal pellets for PCR assay were collected directly from each mouse. Mice were lifted by the base of the tail and allowed to grasp the WBL while a sterile 1.5-mL microfuge tube was held below the anus to allow feces to fall directly into the tube. If the mouse did not defecate within a 30-s period, it was returned to the cage and allowed to rest for at least 2 min before collection was reattempted until a sample was produced.

### Cm PCR

DNA and RNA were copurified from feces or cage swab samples using the Qiagen DNeasy 96 blood and tissue kit (Qiagen, Hilden, Germany). Nucleic acid extraction was performed using the manufacturer’s recommended protocol, “Purification of Total DNA from Animal Tissues”, with the following buffer volume modifications: 300µL of Buffer ATL + Proteinase K, 600µL of buffer AL + EtOH, and 600µL of lysate were added to individual wells of the extraction plate. Washes were performed with 600µL of buffers AW1 and AW2. Final elution volume was 150µL of buffer AE.

A probe-based PCR assay was designed using IDT’s PrimerQuest Tool (IDT, Coralville, IA) based on the 16s rRNA sequence of *Chlamydia muridarum*, Nigg strain (Accession NR_074982.1, located in the National Center for Biotechnology Information database). Primer and Probe sequences generated from the PrimerQuest Tool were checked for specificity using NCBI’s BLAST (Basic Local Alignment Search Tool). The probe was labeled with FAM and quenched with ZEN and Iowa Black FQ (IDT, Coralville, IA). Primer names, followed by associated sequences are as follows: *C. muridarum*_For (GTGATGAAGGCTCTAGGGTTG); *C. muridarum*_Rev (GAGTTAGCCGGTGCTTCTTTA), *C. muridarum*_Probe (TACCCGTTGGATTTGAGCGTACCA).

The Cm PCR assay was validated by generating a standard curve. Positive PCR amplicons derived from known Cm positive fecal samples were combined to a volume of 25µl and purified by pipetting using Diffinity Rapid Tips (Diffinity Genomics, West Chester, PA). Five, 10-fold serial dilutions were run in triplicate producing the following values calculated by Bio-Rad CFX Maestro Software (Bio-Rad, Hercules, CA): E=97.6%, R^2=.995, Slope=-3.381, y-intercept=41.781. To estimate copy numbers, serial dilutions were quantified using a Qubit 2.0 Fluorometer and Qubit dsDNA Quantitation, High Sensitivity Assay (Thermo Fisher Scientific). The concentration values were calculated using the Thermo Fisher DNA Copy Number and Dilution Calculator with default settings for molar mass per base pair, a custom DNA fragment length of 106 bp and a stock concentration of 0.256 ng/µl. From these values, we calculated the Cm PCR assay has a detection limit of 3.36 copies.

Real time (qPCR) PCR was carried out using a BioRad CFX machine (Bio-Rad, Hercules, CA). Reactions were run using Qiagen’s QuantiNova Probe PCR Kit (Qiagen, Hilden, Germany) using the kit’s recommended concentrations and cycling conditions. Final Concentration: 1x QuantiNova Master mix, 0.4 µM Primers, and 0.2 µM FAM labeled Probe. Cycling: 95°C 2 min, followed by 40 cycles of 95°C 5 sec, 60°C 30 sec.

All reactions were run in duplicate by loading 5µl of template DNA to 15µl of the reaction mixture. A positive control and negative, no-template control were included in each run. The positive control was a purified PCR amplicon diluted to produce consistent values of Ct 28. A 16s universal bacterial PCR assay using primers 27_Forward and 1492_Reverse was run on all samples to check for DNA extraction and inhibitors. Samples were considered positive if both replicates had similar values and produced a Ct value of less than 40. A sample was called negative if no amplification from the Cm qPCR assay was detected and a positive amplification from the 16s assay was detected.

### Statistical analysis

Analysis of cage wash temperature through descriptive statistics (mean and standard deviation) was performed using statistics software (Graph Pad Prism 9.1.0, La Jolla, CA). Tunnel washer data are presented as mean ± standard deviation (SD).

## Results

### Intercage transmission study

No transmission of Cm to the pair-housed SW mice occurred. All samples collected from each of the 12 pairs of mice at the 0, 14-, 30-, 90-, and 180-day timepoints were negative for Cm by PCR. Eleven of 12 racks remained positive for Cm by PCR at the end of the study.

### Cage wash study

Figure 2 provides the data logger temperature data for each tunnel washer run (n = 4). A temperature of at least 82°C (180°F) was recorded during each tunnel washer run. The mean maximum temperature for the 4 runs was 84 ± 1°C (182 ± 2°F). The mean amount of time recorded at greater than or equal to 82°C (180°F) was 16 ± 9 s. for the 4 runs.

All 120 (100%) soiled cages (n = 120) were PCR positive for Cm (Table 1) before subsequent processing. None of the cages in the WA or TW group were PCR positive for Cm after processing (Table 1). All pooled fecal samples from SW and NSG mouse pairs in the WA, TW or BR group were PCR negative when tested for Cm 4 weeks after final exposure (Table 1).

**Table 1.** PCR results from cage swabs and test animal feces. “Before” = swabs collected prior to treatment with Tunnel Washer and/or Autoclave. “After” = swabs collected after treatment of positive pretreatment cages. “Feces” = Tac:SW and NSG pairs pooled fecal PCR results at the final collection. All data displayed as positive results out of total samples collected.

| *Treatment Group* | *Timepoint* | *Cm PCR*<br>*result* |
| --- | --- | --- |
| *WA* | **Before** | 100%<br>(40/40) |
|  | **After** | 0% (0/40) |
|  | **Feces** | 0% (0/10) |
| *TW* | **Before** | 100%<br>(40/40) |
|  | **After** | 0% (0/40) |
|  | **Feces** | 0% (0/10) |
| *BR* | **Before** | 100%<br>(40/40) |
|  | **Feces** | 0% (0/10) |

## Discussion

The objective of this study was to assess potential factors that could have contributed to the widespread Cm prevalence within academic research mouse colonies. While our findings offer valuable insights, the factors evaluated do not appear to be contributors to Cm’s prevalence. Our study examined the potential for cage-to-cage transmission resulting from barrier cage changing techniques and commonly used cage sanitization practices to provide insights into Cm’s transmission dynamics within research mouse colonies. Cm transmission did not occur during routine cage changing when employing the strict microisolator cage change techniques practiced in our facility. Perhaps if less stringent husbandry techniques were followed, the risk of transmission would have been higher. Furthermore, our assessment of cage sanitization demonstrated that cages previously housing Cm positive mice and soiled bedding do not serve as vectors for Cm transmission to either immunodeficient or immunocompetent research mice, irrespective of whether the cages were subject to mechanical washing with or without subsequent sterilization in an autoclave. The ubiquitous use of microisolation caging and the associated cage changing techniques has substantially mitigated the spread of numerous adventitious murine pathogens and appears to be effective in containing intercage dissemination of Cm.^10^ Therefore, the question remains as to the factors that have contributed to the relatively high prevalence of this bacterium in research mouse colonies across the globe.^1,26^

The primary mode of transmission of Chlamydia in most species studied is via the fecal-oral route, facilitated by the ingestion of metabolically inactive elementary bodies (EB).^5^ All chlamydial species exhibit a biphasic life cycle, composed of a metabolically inert infectious EB and a metabolically active reticulate body (RB).^15,17,31^ When the EB is internalized by a phagosome in the host mucosal cell, an inclusion body is created.^15,31^ There the EB differentiates into a RB and begins to replicate until cellular lysis or extrusion of newly generated infectious EBs occurs. Released EBs infect nearby cells or are excreted and infect naive hosts.^15,31,34^ *Chlamydial* EB’s typically have a diameter of 0.25 to 0.35 μm and many species, including *C. pneumoniae, C. psittaci, C. suis, C. pecorum, C. felis*, and *C. abortus*, transmit through aerosolized droplets.^3,9,11,27,30,33,36,37^ It is unclear as to whether Cm associated pulmonary disease follows gastrointestinal tract infection and replication, or can Cm be transmitted and infect the lungs directly via inhalation, e.g., as mice root in Cm-contaminated bedding. Disease transmission via aerosolization is categorized by type: droplet transmission, which involves particles larger than 5μm that settle to the ground within 1m of their generation, or airborne transmission, which involves particles smaller than 5μm that can remain suspended in the air for extended periods.^19^ Particles smaller than 10μm are more likely to reach the lower respiratory tract and induce infection.^19^ Notably, *C. pneumoniae* has been shown to transmit to mouse lungs through aerosolized particles with a size of 5μm.^16^ Given that Cm EBs are of a similar size of 0.2 to 0.3 μm, it is plausible that aerosol transmission can occur, however we did not see evidence of it in the current study. ^14,19,27,35,37^

In this regard, Cm appears to differ significantly from *C. bovis*, a relatively common pathogen of immunodeficient mice.^4,6,8,26^ In contrast to Cm, *C. bovis* can be transmitted between cages when following strict microisolator cage changing technique in a biological safety cabinet as it is easily aerosolized.^6^ While IVCs effectively prevented *C. bovis* contamination of cages when maintained on the ventilated rack in a room with high levels of aerosolized *C. bovis*, the contamination occurred when the cage was manipulated during cage changing.^6^ While not evaluated in this study, the caging system employed, which was the same used by Burr *et al*., would be expected to exclude infectious Cm EBs if they were aerosolized.^6^ Based on the lack of transmission during cage change in this study, Cm EBs are either not easily aerosolized during cage change or, if they are, the number of aerosolized EBs is below the minimum infectious dose.^39^ Yeruva et al (2013) concluded that the ID_50_ for Cm was less than 100 inclusion forming units.^39^ The transmissibility of *C. bovis* is attributed to its small size, extracellular nature, lipophilicity and keratotropic properties, facilitating its aerosolization on skin flakes.^4,6^ Additionally, *C. bovis* has been shown to survive in the environment for over 30 days.^6^ Collectively, these features contribute to its infectivity.^4,6^ In contrast, *Chlamydial* species, including Cm, lack the environmental persistence and transmissibility observed by *C. bovis* in vivaria.^20,28,38^ *C. trachomatis* and *C. pneumoniae* remain infectious in the environment for up to 240 and 180 minutes, respectively, a considerably shorter period than observed with *C. bovis*.^20,28,38^

We also assessed whether inadequate cage sanitation procedures could contribute to inadvertent transmission of Cm via fomites. Chlamydia are susceptible to both disinfectants and heat.^12,38^ *C. trachomatis* can be fully inactivated by 0.25% bleach, 70% ethanol, 7.5% hydrogen peroxide or 0.2% peracetic acid, and complete inactivation occurs at 131°F (55°C) for a minimum of 10 minutes.^12^ We speculated that Cm could have similar inactivation requirements and had been inadvertently transmitted within our program via contaminated caging as our standard practice for rodent cage sanitation involves washing cages in a mechanical washer reaching 180°F (82.2°C) without chemicals and without subsequent autoclaving, as both have been associated with accelerated thermoplastic degradation leading to the leaching of monomeric components, many of which are recognized endocrine disruptors.^21^ Our findings demonstrate that Cm is not easily transmitted via contaminated caging as neither immunocompetent nor highly immunocompromised mice became infected after housing in untreated cages which had housed mice shedding Cm and were Cm PCR-positive prior to housing, suggesting the presence of free EBs and/or Cm nucleic acid adhering to the surfaces of the caging. This result may reflect that either the nucleic acid was not infectious, or the amount present was below the minimal infectious dose.^39^ While appearing to be irrelevant, Cm nucleic acid was effectively removed by mechanical washing.

Importantly, although soiled bedding-free, Cm-contaminated caging did not result in transmission of Cm in this study, evidence from dirty bedding sentinels enrolled in our colony health monitoring program supports transmission via soiled bedding.^26,29^ Therefore, Cm could potentially be transmitted between cages in close proximity to one another through dispersal of contaminated bedding and fecal material if mice are housed in open-top caging, or between mice, if behavioral and surgical equipment is not adequately cleaned between uses. Conversely, the potential for transmission among microisolation cages is low when appropriate husbandry practices are employed. Therefore, treatment or rederivation of select mouse strains to eliminate Cm would likely not be compromised through reinfection by Cm-positive lines maintained in the same room or facility if direct contact with Cm positive animals or their feces is prevented.

Given that neither of the factors assessed in the current study likely contribute to Cm transmission, the question as to how Cm became so prevalent remains unanswered. Intra and interinstitutional sharing and breeding of research mice emerges as the most probable culprit. Drawing from data in our program’s databases, we sought data that would support this hypothesis. We did not begin testing for Cm during quarantine or within our colony health monitoring program until 2021, allowing Cm to propagate unchecked for perhaps decades. A total of 14,378 research mice were imported from academic institutions for use by investigators in our program between November 2013 to December 2019. These mice were primarily unique GEM strains that were dispersed to 184 PI’s. These quarantined mice originated from 280 domestic and international institutions, representing a total of 685 PIs at these institutions. Using the current Cm positivity rate of 33% of recent importations, we could have inadvertently introduced 4,313 positive mice into our colonies over this 6-year period. To illustrate the potential for intrainstitutional spread of Cm, 1042 intrainstitutional transfers of 1 to 6 cages of mice occurred between July 2015 and December 2019 in our program which reflects the sharing of 4000 and 32,000 mice between laboratories over this 4.5-year period. We also estimated the potential for Cm-positive mice to have been distributed to other institutions. We distributed mice from 90 rodent housing rooms to 169 receiving institutions from January 2018 to December 2019, and 89.8% of these housing rooms are currently Cm positive. If our program is representative of murine import and export activity, and intrainstitutional sharing of mice at other major biomedical research programs, we believe that the unknowing distribution of Cm-infected mice was sufficient to have resulted in the global and intrainstitutional prevalence currently seen, given that most institutions use a test and exclusion importation strategy.^26^ As vendor maintained mouse colonies have been rederived and maintained in barriers and have consistently tested negative for Cm, we do not believe they are a likely source of infection. While sharing infected mice could have led to the currently observed prevalence, it does not shed light into how and how many times Cm was initially introduced into one or more colonies across the globe to eventually lead up to the current levels of infection. While only speculation, we believe it could reflect “escape” of an experimentally used isolate since we have not detected Cm in wild mice which we tested.

In conclusion, this study aimed to gain further insight into the factors that could have contributed to the prevalence of Cm in academic research colonies. We concluded that neither cage change technique nor cage sanitation practices contributed to the spread of Cm within our program. We suspect Cm transmission requires animal-to-animal or soiled bedding-to-animal contact suggesting that elimination of Cm may be straightforward with the use of an efficacious antimicrobial agent. Studies are ongoing to identify such a therapeutic. Given the potential for Cm infections to cause various biological effects across mouse strains, including mortality in immunodeficient strains, we suggest that Cm be treated as an ‘excluded’ agent. Therefore, it is critical to implement institutional biosecurity protocols to prevent its introduction and facilitate its eradication from infected colonies.

## Abbreviations

Cm: *Chlamydia muridarum*
GEM: genetically engineered mouse
NSG: NOD.Cg-*Prkdc*^*scid*^ *Il2rg*^*tm1Wjl*^/SzJ
SW: Tac:SW
ACS: animal changing station
WBL: wire bar lid
IVC: individually ventilated cage
TW: tunnel wash treatment group
WA: wash-then-autoclaved treatment group

## Acknowledgements

MSK Core Facilities are supported by MSK’s NCI Cancer Center Support Grant P30 CA008748. The authors would like to thank Amanda Ritter for study design guidance, Abigail Michelson and Seth Lieberman for support during data collection, Simona Bekker and John D’Allara for their technical assistance during sample handling and shipment, Fabricio Munoz and Marcia Lewis for their assistance with breeding of the NSG mice used in this study, Stanley Fairley and Jose Monterosa for their assistance with the operation of the tunnel washer and autoclave, Jack Palillo for statistical guidance, and James Fahey for diagnostic oversight.

## References

1. Albers TM, Henderson KS, Mulder GB, Shek WR. 2023. Pathogen Prevalence Estimates and Diagnostic Methodology Trends in Laboratory Mice and Rats from 2003 to 2020. J Am Assoc Lab Anim Sci 62:229–242. 10.30802/AALAS-JAALAS-22-000097

2. American Veterinary Medical Association. 2020. AVMA guidelines for the euthanasia of animals: 2020 edition. American Veterinary Medical Association :1–111.

3. Balsamo G, Maxted AM, Midla JW, Murphy JM, Wohrle R, Edling TM, Fish PH, Flammer K, Hyde D, Kutty PK, et al. 2017. Compendium of Measures to Control Chlamydia psittaci Infection Among Humans (Psittacosis) and Pet Birds (Avian Chlamydiosis), 2017. J Avian Med Surg 31:262–282. 10.1647/217-265

4. Besch-Williford C, Franklin C. 2007. Aerobic Gram-Positive Organisms, p 399–401. In: Fox J, Davisson M, Quimby F, Barthold S, Newcomer C, Smith A, editors. The mouse in biomedical research. 2nd ed. Burlington: Elsevier.

5. Boleti H. 2007. Chlamydia Infections, p 1–9. In: Enna S, Bylund David B, editors. xPharm: The Comprehensive Pharmacology Reference. Amsterdam: Elsevier Inc.

6. Burr H, Wolf F, Lipman N. 2012. Corynebacterium bovis: epizootiologic features and environmental contamination in an enzootically infected rodent room. J Am Assoc Lab Anim Sci 51:189–198.

7. Carlson AL, Floyd RJ, Ricart Arbona RJ, Henderson KS, Perkins C, Lipman NS. 2022. Assessing Elimination of Mouse Kidney Parvovirus from Cages by Mechanical Washing. J Am Assoc Lab Anim Sci 61:61–66. 10.30802/AALAS-JAALAS-21-000096

8. Clifford CB, Walton BJ, Reed TH, Coyle MB, White WJ, Amyx HL. 1995. Hyperkeratosis in Athymic Nude Mice caused by a Coryneform bacterium: Microbiology, Transmission, Clinical Signs and Pathology. Lab Anim Sci 45:131–139.

9. Cole EC. 1990. The Chlamydiae: Infectious Aerosols in Indoor Environments, p 99–113. In: Morey P, Feeley J, Otten JA, editors. Biological Contaminants in Indoor Environments. Baltimore: ATSM International.

10. Compton SR, Homberger FR, MacArthur Clark J. 2004. Microbiological monitoring in individually ventilated cage systems. Lab Anim 33:36–41. 10.1038/laban1104-36

11. Corsaro D, Venditti D. 2004. Emerging chlamydial infections. Crit Rev Microbiol 30:75–106. 10.1080/10408410490435106

12. Coulon C, Eterpi M, Greub G, Collignon A, McDonnell G, Thomas V. 2012. Amoebal host range, host-free survival and disinfection susceptibility of environmental Chlamydiae as compared to Chlamydia trachomatis. FEMS Immunol Med Microbiol 64:364–373. 10.1111/j.1574-695X.2011.00919.x

13. Dochez A, Mills K, Mulliken B. 1937. A Virus Disease of Swiss Mice Transmissible by Intranasal Inoculation. Proc Soc Exp Biol Med 36:683–686. 10.3181/00379727-36-9357

14. Droogenbroeck C Van, Risseghem M Van, Braeckman L, Vanrompay D. 2009. Evaluation of bioaerosol sampling techniques for the detection of Chlamydophila psittaci in contaminated air. Vet Microbiol 135:31–37. 10.1016/j.vetmic.2008.09.042

15. Elwell C, Mirrashidi K, Engel J. 2016. Chlamydia cell biology and pathogenesis. Nat Rev Microbiol 14:385–400. 10.1038/nrmicro.2016.30

16. Falsey AR, Walsh EE. 1993. Transmission of Chlamydia pneumoniae. J Infect Dis 168:493–6. 10.1093/infdis/168.2.493

17. Gitsels A, Sanders N, Vanrompay D. 2019. Chlamydial Infection From Outside to Inside. Front Microbiol 10:1–27.

18. Gordon FB, Freeman G, Glampit J. 1938. A pneumonia-producing filtrable agent from stock mice. Proc Soc Exp Biol Med 39:450–453.

19. Gralton J, Tovey E, McLaws M-L, Rawlinson WD. 2011. The role of particle size in aerosolised pathogen transmission: A review. J Infect 62:1–13. 10.1016/j.jinf.2010.11.010

20. Hammerschlag MR, Kohlhoff SA, Gaydos CA. 2015. Chlamydia pneumoniae, p 2174–2182. In: Bennett JE, Dolin R, Blaser MJ, editors. Mandell, Douglas, and Bennett’s Principles and Practice of Infectious Diseases. 2nd ed. New York: Saunders.

21. Howdeshell KL, Peterman PH, Judy BM, Taylor JA, Orazio CE, Ruhlen RL, Saal FS vom, Welshons W V. 2003. Bisphenol A is released from used polycarbonate animal cages into water at room temperature. Environ Health Perspect 111:1180–1187. 10.1289/ehp.5993

22. Institute for Laboratory Animal Research. 2011. Guide for the Care and Use of Laboratory Animals. 8th ed. Washington, D.C.: National Academies Press.

23. Lipman NS, Homberger FR. 2003. Rodent quality assurance testing: use of sentinel animal systems. Lab Anim 32:36–43. 10.1038/laban0503-36

24. Mishkin N, Carrasco SE, Miranda IC, Cheleuitte-Nieves C, Ricart Arbona RJ, Lipman N S. 2024. Chlamydia muridarum Associated Pulmonary and Urogenital Disease and Pathology in a Colony of Enzootically-Infected IL12rb2 deficient and Stat1 knockout Mice. Comp Med :Online ahead of print. 10.30802/AALAS-CM-24-000002

25. Mishkin N, Palillo MB, Otis C, Longhini AL, Gardner R, Ricart Arbona RJ, Carrasco SE, Palillo JA, Joseph A, Sonnenberg GF, Lipman NS. 2023. Chlamydia muridarum Modulates Splenic Monocyte and T-cell Response and Induces Sustained Intestinal T-cell and ILC3 Responses in Inbred and Outbred Mice. Abstracts of Scientific Presentations presented at the 2023 AALAS National Meeting, Salt Lake City, Utah. J Am Assoc Lab Anim Sci 62:577.

26. Mishkin N, Ricart Arbona RJ, Carrasco SE, Lawton S, Henderson KS, Momtsios P, Sigar IM, Ramsey KH, Cheleuitte-Nieves C, Monette S, Lipman NS. 2022. Reemergence of the Murine Bacterial Pathogen Chlamydia muridarum in Research Mouse Colonies. Comp Med 72:230–242. 10.30802/AALAS-CM-22-000045

27. Miyashita N, Matsumoto A. 2005. Morphology of Chlamydia pneumoniae, p 11–26. In: Friedman H, Yamamoto Y, Bendinelli M, editors. Chlamydia pneumoniae Infection and Disease. New York: Kluwer Academic/Plenum Publishers.

28. Novak K, Kowalski R, Karenchak L, Gordon Y. 1995. Chlamydia trachomatis can be transmitted by a nonporous plastic surface in vitro. Cornea 14:523–536.

29. Palillo M, Lipman N, Ricart Arbona R. 2022. Effectiveness of Various Antibiotics for Treating Chlamydia muridarum-infected mice. Abstracts of Scientific Presentations presented at the 2022 AALAS National Meeting, Louisville Kentucky. J Am Assoc Lab Anim Sci 61:506–589.

30. Puysseleyr K De, Puysseleyr L De, Dhondt H, Geens T, Braeckman L, Morré SA, Cox E, Vanrompay D. 2014. Evaluation of the presence and zoonotic transmission of Chlamydia suis in a pig slaughterhouse. BMC Infect Dis 14:1–6. 10.1186/s12879-014-0560-x

31. Rank R. 2007. Chlamydial Diseases, p 326–344. In: Fox JG, Davisson MT, Quimby FW, Barthold SW, Newcomer CE, Smith AL, editors. The mouse in biomedical research. Vol. 2. Burlington: Elsevier.

32. Ritter AC, Ricart Arbona RJ, Mourino AJ, Palillo MB, Aydin M, Fahey JR, Lipman NS. 2023. Mechanical Washing Prevents Transmission of Bacterial, Viral, and Protozoal Murine Pathogens from Cages. J Am Assoc Lab Anim Sci 62:131–138. 10.30802/AALAS-JAALAS-22-000105

33. Sachse K, Grossmann E, Berndt A, Schütt C, Henning K, Theegarten D, Anhenn O, Reinhold P. 2004. Respiratory chlamydial infection based on experimental aerosol challenge of pigs with Chlamydia suis. Comp Immunol Microbiol Infect Dis 27:7–23. 10.1016/S0147-9571(02)00079-6

34. Scidmore MA. 2009. Chlamydia, p 74–86. In: Schaechter M, editor. Encyclopedia of Microbiology. 3e ed. Internet: Academic Press.

35. St Jean SC, Ricart Arbona RJ, Mishkin N, Monette S, Wipf JRK, Henderson KS, Cheleuitte-Nieves C, Lipman NS, Carrasco SE. 2024. Chlamydia muridarum infection causes bronchointerstitial pneumonia in NOD.Cg-PrkdcscidIl2rgtm1Wjl/SzJ (NSG) mice. Vet Pathol 61:145–156. 10.1177/03009858231183907

36. Theunissen HJ, Lemmens-den Toom NA, Burggraaf A, Stolz E, Michel MF. 1993. Influence of temperature and relative humidity on the survival of Chlamydia pneumoniae in aerosols. Appl Environ Microbiol 59:2589–2593. 10.1128/aem.59.8.2589-2593.1993

37. Theunissen JH. 1993. Chlamydia, physical characteristics and diagnostic aspects. Thesis: Erasmus University Rotterdam: 9–22.

38. Verkooyen RP, Harreveld S, Ahmad Mousavi Joulandan S, Diepersloot RJ, Verbrugh H. 1995. Survival of Chlamydia pneumoniae following contact with various surfaces. Clin Microbiol Infect 1:114–118.

39. Yeruva L, Spencer N, Bowlin AK, Wang Y, Rank RG. 2013. Chlamydial infection of the gastrointestinal tract: A reservoir for persistent infection. Pathog Dis 68:88–95. 10.1111/2049-632X.12052

